# Oxytocin excites BNST interneurons and inhibits BNST output neurons to the central amygdala

**DOI:** 10.1101/2020.06.24.169466

**Authors:** Walter Francesconi, Fulvia Berton, Valentina Olivera-Pasilio, Joanna Dabrowska

## Abstract

The dorsolateral bed nucleus of the stria terminalis (BNST_DL_) has high expression of oxytocin (OT) receptors (OTR), which were shown to facilitate cued fear. However, the role of OTR in the modulation of BNST_DL_ activity remains elusive. BNST_DL_ contains GABA-ergic neurons classified based on intrinsic membrane properties into three types. Using *in vitro* patch-clamp recordings in male rats, we demonstrate that OT selectively excites and increases spontaneous firing rate of Type I BNST_DL_ neurons. As a consequence, OT increases the frequency, but not amplitude, of spontaneous inhibitory post-synaptic currents (sIPSCs) selectively in Type II neurons, an effect abolished by OTR antagonist or tetrodotoxin, and reduces spontaneous firing rate in these neurons. These results suggest an indirect effect of OT in Type II neurons, which is mediated via OT-induced increase in firing of Type I interneurons. As Type II BNST_DL_ neurons were shown projecting to the central amygdala (CeA), we also recorded from retrogradelly labeled BNST→CeA neurons and we show that OT increases the frequency of sIPSC in these Type II BNST→CeA output neurons. In contrast, in Type III neurons, OT reduces the amplitude, but not frequency, of both sIPSCs and evoked IPSCs via a postsynaptic mechanism without changing their intrinsic excitability. We present a model of fine-tuned modulation of BNST_DL_ activity by OT, which selectively excites BNST_DL_ interneurons and inhibits Type II BNST→CeA output neurons. These results suggest that OTR in the BNST might facilitate cued fear by inhibiting the BNST→CeA neurons.

**Highlights:**

- Oxytocin directly excites and increases spontaneous firing of Type I BNST interneurons
- Oxytocin indirectly inhibits Type II BNST neurons
- Oxytocin inhibits Type II BNST output neurons to the central amygdala (BNST→CeA)
- Oxytocin reduces GABA-ergic transmission in Type III BNST neurons
- Oxytocin might facilitate cued fear by inhibiting the Type II BNST→CeA neurons

**Graphical abstract:** 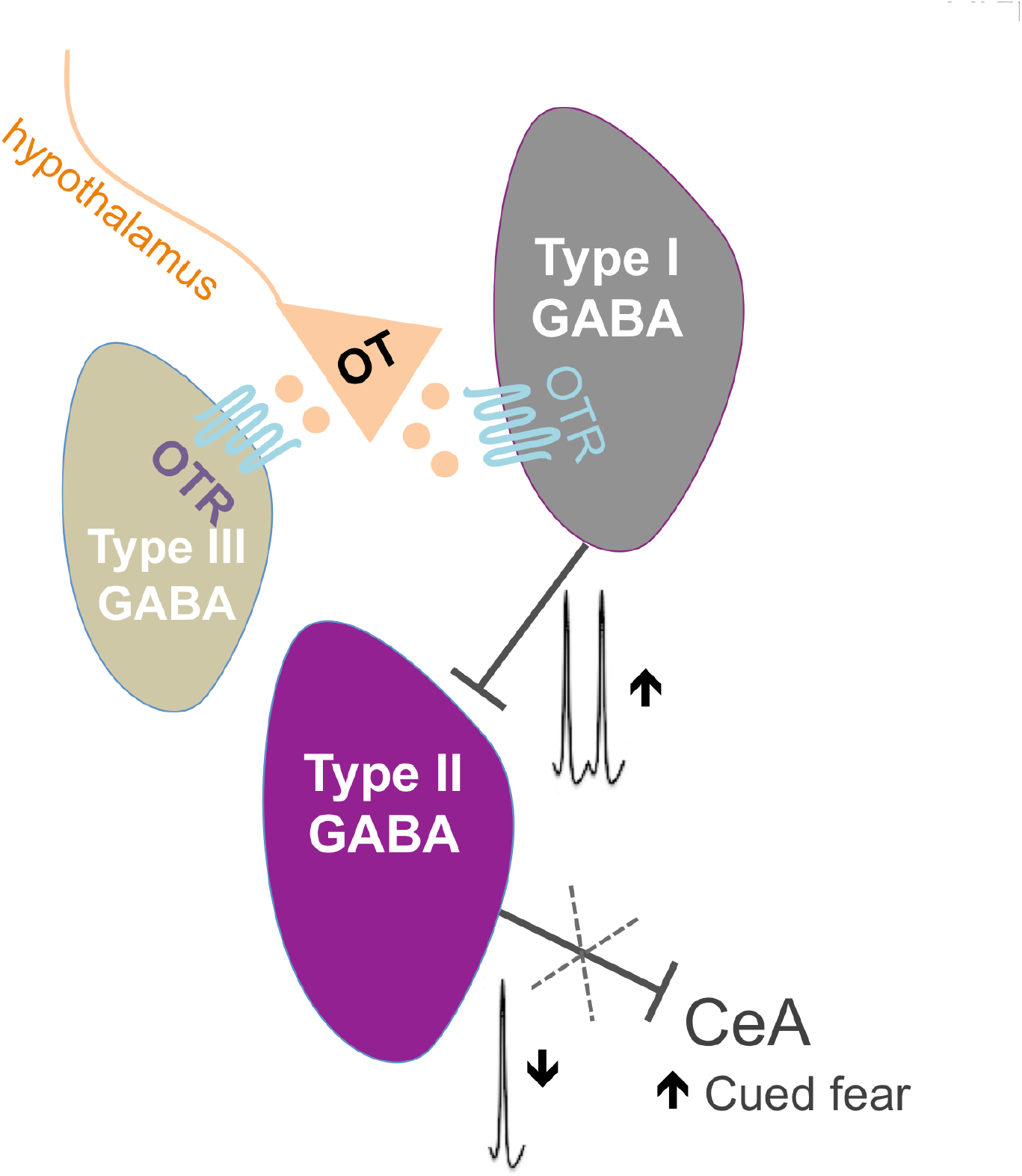

## 1. Introduction

Oxytocin (OT) is a nine amino acid peptide serving as a hypothalamic hormone, which regulates uterine contractions during labor, milk ejection reflex during lactation (Caldeyro-Barcia and Poseiro, 1959; Nickerson et al., 1954), as well as water-electrolyte balance (Han et al., 1993; Verbalis et al., 1993). As a neuromodulator in the central nervous system, OT facilitates social behaviors (Bosch and Young, 2018; Pedersen et al., 1992; Tabbaa et al., 2016), regulates stress response, stress-coping behaviors, and fear and anxiety-like behaviors, for review see (Janeček and Dabrowska, 2019; Neumann and Slattery, 2016; Olivera-Pasilio and Dabrowska, 2020). Although OT has been shown to reduce fear responses primarily in the central amygdala (CeA) (Knobloch et al., 2012; Terburg et al., 2018; Viviani et al., 2011), we recently showed that oxytocin receptor (OTR) transmission in the dorsolateral bed nucleus of the stria terminalis (BNST_DL_) facilitates the formation of cued fear and improves fear discrimination to signaled vs. un-signaled threats (Martinon et al., 2019; Moaddab and Dabrowska, 2017), for review see (Olivera-Pasilio and Dabrowska, 2020).

On a cellular level, OT has shown robust modulatory influence on the inhibitory neurocircuitry of the hippocampus (Harden and Frazier, 2016; Maniezzi et al., 2019; Owen et al., 2013; Tirko et al., 2018), auditory cortex (Marlin et al., 2015), ventral tegmental area, VTA, (Hung et al., 2017; Xiao et al., 2017), lateral amygdala (Crane et al., 2020), and the CeA (Huber et al., 2005; Knobloch et al., 2012; Terburg et al., 2018; Viviani et al., 2011). Notably, although the BNST has one of the highest expression levels of OTR in the rodent brain (Dabrowska et al., 2011; Dumais et al., 2013; Tribollet et al., 1992; Veinante and Freund-Mercier, 1997) and receives OT inputs from hypothalamic nuclei (Dabrowska et al., 2011; Knobloch et al., 2012; Martinon et al., 2019), the effects of OT on excitability and synaptic transmission of BNST neurons remain elusive. Studies in the 90’s from the Ingram group using electrophysiological approaches have shown that OT increases the spontaneous firing frequency in a subpopulation of BNST neurons in lactating female rats (Ingram et al., 1990; Ingram and Moos, 1992; Wilson et al., 2005), but their type was not known.

BNST_DL_ neurons are GABA-ergic (Cullinan et al., 1993; Dabrowska et al., 2013a; Sun and Cassell, 1993), and functionally interconnected via a dense intrinsic inhibitory network (Xu et al., 2016). As originally described by Rainnie and colleagues, BNST_DL_ neurons can be classified into three major neuron types (I-III) based on their electrophysiological phenotype defined as a unique pattern of voltage deflection in response to transient depolarizing and hyperpolarizing current injections (Daniel et al., 2017; Hammack et al., 2007) and/or based on a single-cell mRNA expression of ion channel subunits (Hazra et al., 2011). Recently, it was shown that both Type II and Type III are output neurons of the BNST_DL_, with major projections to the CeA (mainly Type II neurons), lateral hypothalamus (LH), and the VTA (mainly Type III neurons) (Yamauchi et al., 2018).

In the current study, using *in vitro* slice patch-clamp recordings in a whole-cell and cell-attached mode, we investigated the effects of OT on intrinsic excitability, inhibitory synaptic transmission and spontaneous firing in Type I-III BNST_DL_ neurons in male Sprague-Dawley rats. We show that OT is a powerful modulator of the intrinsic inhibitory network such as it selectively increases the intrinsic excitability and the frequency of spontaneous firing of regular spiking, Type I neurons, which leads to increased inhibitory synaptic transmission (spontaneous Inhibitory Postsynaptic Currents, sIPSCs) and reduced spontaneous firing rate of Type II BNST_DL_ neurons. We also combined injections of a fluorescent retrograde tracer into the CeA with electrophysiological recordings from fluorescent BNST_DL_ neurons and we confirm that all recorded CeA-projecting neurons (BNST→CeA) are Type II neurons. Notably, we show that OT increases the sIPSCs frequency in these Type II BNST→CeA projection neurons. Finally, we demonstrate that OT reduces the amplitude of both spontaneous and evoked IPSCs selectively in Type III neurons via an OTR-dependent, postsynaptic mechanism. These results demonstrate that OT directly excites Type I BNST_DL_ interneurons, whereas it indirectly inhibits the Type II BNST→CeA projection neurons.

## 2. Material and methods

### 2.1. Animals

Male Sprague-Dawley rats (Envigo, Chicago, IL; 240-300 g) were housed in groups of three on a 12 h light/dark cycle (light 7 a.m. to 7 p.m.) with free access to water and food. Rats were habituated to this environment for one week before the experiments began. Experiments were performed in accordance with the guidelines of the NIH and approved by the Animal Care and Use Committee at RFUMS. A total of 153 rats were used for the experiments. The effects of OT on membrane properties were tested in 14 Type I neurons (from 10 rats), 13 Type II (12 rats), and 9 Type III neurons (8 rats). The effect of OT on evoked firing was tested on 12 Type I (11 rats), 8 Type II (7 rats), and 9 Type III neurons (8 rats). The effect of OT on the spontaneous firing was tested in 9 Type I (8 rats) and 19 Type II (16 rats). To study the effect of OT on sIPSCs, we recorded 14 Type II (12 rats), 6 Type I (5 rats), and 8 Type III neurons (8 rats). We recorded 6 Type II neurons (4 rats) in experiment with OTR antagonist, and 6 Type II neurons (5 rats) in TTX experiment. The effect of OT on the evoked IPSCs was tested in 5 Type I (5 rats), 8 Type II (8 rats), and 6 Type III (5 rats) neurons. We recorded 5 Type III (4 rats) neurons in experiments with OTR antagonist and 6 Type III (4 rats) with GABA-B antagonist. We recorded 13 retrogradelly labeled Type II BNST→CeA neurons from 11 rats and we recorded sIPSCs after application of OT in 5 of these Type II BNST→CeA neurons (from 5 rats).

### 2.2 Slice preparation for electrophysiology

Rats were anaesthetized with Isoflurane USP (Patterson Veterinary, Greeley, CO, USA) before decapitation. Brains were quickly removed and coronal 300 µm slices containing the BNST_DL_ were prepared in ice-cold cutting solution (saturated with 95% O_2_/ 5% CO_2_) containing in mM: NaCl 130, KCl 3.5, NaHCO_3_ 30, KH_2_PO_4_ 1.1, CaCl_2_·2H_2_O 1, D-glucose 10, MgCl_2_ 6H_2_O 6, Kynurenate 2, pH 7.4, 290-300 mOsm. The slices were prepared using a Leica vibratome (VT1200; Leica, Wetzlar, Germany), then incubated for 1 hr at 34°C in the cutting solution and subsequently transferred to room temperature in artificial cerebrospinal fluid (aCSF) containing in mM: NaCl 130, KCl 3.5, NaHCO_3_ 30, KH_2_PO_4_ 1.1, CaCl_2_·2H_2_O 2.5, D-glucose 10, and MgCl_2_·6H_2_O 1.3, Ascorbate 0.4, Thiourea 0.8, Na Pyruvate 2, pH 7.4, with osmolarity of 290-300 mOsm, continuously oxygenated with 95% O_2_/ 5% CO_2_. Next, the slices were transferred to a submerged recording chamber and perfused with aCSF at 33°C ± 1°C saturated with 95% O_2_/ 5% CO_2_ at a flow rate of 2-3 ml/min. BNST_DL_ neurons were visualized using infrared differential interference contrast (IR-DIC) optic with an Upright microscope (Scientifica SliceScope Pro 1000, Clarksburg, NJ, USA).

### 2.3. Patch-clamp recordings

Whole-cell patch-clamp recordings were performed from BNST_DL_ neurons, primarily in the oval nucleus of the BNST (BNST_OV_), using glass pipette (4-8 MΩ) pulled from thick-walled borosilicate glass capillaries with a micropipette puller (Model P-97; Sutter Instrument, Novato, CA, USA) as before (Dabrowska et al., 2013b; Francesconi et al., 2009). To study the effects of OT (0.2 μM concentration based on (Crane et al., 2020; Xiao et al., 2017), oxytocin acetate salt, # H-2510, Bachem, Switzerland) on intrinsic membrane properties and firing activity the patch pipettes were filled with potassium gluconate internal solution containing in mM: K-gluconate 135, KCl 2, MgCl_2_·6H_2_O 3, HEPES 10, Na-phosphocreatine 5, ATP-K 2, GTP-Na 0.2, pH 7.3 adjusted with KOH, 300-305 mOsm. Intrinsic membrane properties experiments were performed in a presence of synaptic transmission blockers: the AMPA receptor antagonist CNQX (10 μM), the NMDA receptor antagonist AP-5 (50 μM), and GABA-A receptor antagonist, picrotoxin (25 μM). To study the effects of OT on the spontaneous inhibitory postsynaptic currents (sIPSCs), the patch pipettes were filled with high chloride internal solution in mM: KOH 110, MgCl_2_· 6H_2_O 2, KCl 50, EGTA 0.2, ATP-Na 2, GTP-Na 0.3, HEPES 10, and Spermine 0.1, the pH was adjusted to 7.3 with D-gluconic acid and osmolarity was 300-305 mOsm. The evoked IPSCs (eIPCSs) were recorded with pipettes filled with K-gluconate solution as described above. In order to isolate GABAergic eIPSCs, the eIPSCs were recorded at the holding voltage (Vh) of +10 mV, which is the reversal potential for the excitatory postsynaptic current. A stimulating pipette filled with aCSF was placed in the Stria Terminalis. For each neuron recorded the stimulus intensity was adjusted to give an eIPSCs of about 1/3 of the maximal amplitude and maintained throughout the experiment. We applied 10 paired pulse stimuli (ISI 50 msec) at the frequency of 0.2 Hz. The peak amplitude of the eIPSC was measured from the averaged responses. The responses were acquired each minute for 10 min before and 12 min after OT application. The paired pulse stimulus (50 ms ISI) was used to calculate the paired pulse ratio (PPR) between the mean amplitude of the second eIPSC divided by the amplitude of the first eIPSC. Here we used the PPR to evaluate a possible presynaptic mechanism of the OT effect on the BNST neurons inhibitory synaptic responses. To further examine a possible presynaptic mechanism due to an activation of presynaptic GABA-B receptors, in a subset of experiments we applied a specific GABA-B receptor antagonist CGP 55845 (2 μM) before OT.

To verify that the effects of OT were mediated by OTRs, selective and potent OTR antagonist d(CH2)5(1), D-Tyr(2), Thr(4), Orn(8), des-Gly-NH2(9)]-Vasotocin trifluoroacetate salt (Manning et al., 2012) (0.4 μM, OTA, V-905 NIMH Chemical Repository) was applied 10 min before OT in all whole-cell experiments. Recordings were performed with a Multiclamp 700B amplifier (Molecular Device, Sunnyvale, CA, USA). Current-clamp and voltage-clamp signals were acquired at 10 kHz with a 16-bit input-output board NI USB-6251 (National Instruments, Austin, TX) using custom MatLab scripts (Math-Work, Natick, MA) written by Dr. Niraj Desai (Desai et al., 2017). All neurons included in the analysis had a seal resistance > 1GΩ and an initial access resistance < 20 MΩ maintained for the entire recording period. Any cells showing a 10% change in the serial resistance were not included in the analysis.

#### 2.3.1. Electrophysiological characterization of BNST_DL_ neurons

At the beginning of the recording sessions in current clamp mode, neurons were electrophysiologically characterized into three types (Type I-III) using current pulses (450 msec in duration) from -350 pA to 80 pA in 5 or 10 pA increments. Based on their characteristic voltage responses (Hammack et al., 2007), three types of neurons were identified in the BNST_DL_. Type I neurons are regular firing and display spike frequency adaptation. These neurons also display moderate voltage sag that indicates the presence of the hyperpolarization-activated cation current (I_h_). As a distinguishing feature of Type II neurons, post-inhibitory spikes are produced in response to preceding negative current steps that are related to the action of the low-threshold Ca^2+^ current. Additionally, Type II neurons display strong voltage sags under hyperpolarizing current pulses that indicate a high level of I_h_ current. Type III neurons differ from the previous two types in several aspects: they exhibit high rheobase and no voltage sag under negative current steps; they display prominent inward rectification that is caused by the activation of the inward rectifying K^+^ current (I_KIR_) at membrane potentials more negative than approximately -50 mV; and start firing after a characteristic slow voltage ramp mediated by the K^+^ delayed current (I_D_).

#### 2.3.2. Cell-attached recordings

The spontaneous firing activity of Type I and II BNST_DL_ neurons was recorded in a cell-attached voltage-clamp mode, using patch pipettes filled with K-gluconate solution as described above, without blockers of the synaptic transmission. To avoid a possible effect of amplifier-generated currents on the spontaneous firing activity during the recording the command potential was carefully adjusted at the value at which no current was flowing from the amplifier through the cell membrane as described before (Perkins, 2006). At the end of the spontaneous firing recording period, a gentle negative pressure was applied to break through the membrane into whole cell configuration and determine the neuron type in a current-clamp mode as described above.

#### 2.3.3. The effect of OT on intrinsic membrane properties and spontaneous firing of Type I-III BNST_DL_ neurons

We investigated the effects of OT on intrinsic membrane properties: resting membrane potential (RMP), input resistance (Rin), rheobase (Rh), threshold of the first action potential (Th) (calculated as the voltage at which the depolarization rate exceeded 5 mV/ms), and latency of the first spike (Lat) evoked at the Rh current. We also investigated spiking ability by measuring the input-output (I-O) relationship before and after OT application. The steady state frequency (SSF) was measured by applying depolarizing pulses (1 sec in duration) of different amplitudes (0-80 pA in 10 pA increments). The SSF was determined as the average of the inverse of the inter-spike intervals (ISI) from all action potentials starting from the second action potential. The I/O curve was built before (pre), during, and after OT application (post).

A variety of afterhyperpolarizations (AHPs) following a single or a train of action potentials have been described to regulate the neuron excitability and spike frequency adaptation (Gu et al., 2008; Lopez de Armentia and Sah, 2004). They come in a temporal sequence of three AHPs: the fast AHP (fAHP) lasting 1-5 msec, the medium AHP (mAHP) lasting 50-200 msec, which is responsible for early spike frequency adaptation during a train of action potentials, and the late AHP (lAHP) lasting several seconds after a train of action potentials (Gu et al., 2005a). We investigated the effect of the OT on the amplitude of fAHP, mAHP and lAHP in all types of the BNST_DL_ neurons. The amplitudes of the fAHP and mAHP following the first action potential evoked at Rh were calculated as the difference from the action potential threshold to the membrane voltage measured after 2-5 msec and 50-100 msec, respectively. The lAHP was measured at 2000 msec after the end of the depolarizing pulse, following generation of five action potentials. As the K^+^ currents underlying these AHPs are considered to regulate spike frequency adaptation (Gu et al., 2008; Lopez de Armentia and Sah, 2004), we measured early and late spike frequency adaptation before and after OT application using a depolarizing pulse of 1 sec. We measured the instantaneous frequency for every action potential interval. The ISI number is the inter-stimulus interval between two consecutive action potentials. We measured the spike frequency adaptation rather than the spike frequency, because the firing of the BNST neurons, as described by Paré and colleagues (Rodríguez-Sierra et al., 2013), is adapting and not regular. This is a characteristic of all types of BNST_DL_ neurons, including Type I (described as regular firing), which also adapt on a long current pulse. We also performed recordings in cell-attached mode in Type I-II neurons and measured the spontaneous firing frequency before and after OT. The spontaneous firing frequency (Hz) was determined as the number of spontaneous spikes for every minute of recording.

#### 2.3.4. The effect of OT on inhibitory synaptic activity of Type I-III BNST_DL_ neurons

To study the effect of OT on spontaneous inhibitory postsynaptic currents (sIPSCs), the events were recorded from Type I-III neurons in a voltage clamp mode at holding potential of -70 mV. To isolate the sIPSCs, the recordings were made in the presence of the AMPA receptor antagonist CNQX (10 μM) and the NMDA receptor antagonist AP-5 (50 μM). The sIPSCs were characterized by their mean frequency, amplitude, and charge (calculated as the area under the curve) using custom scripts in MatLab. To reach an adequate number of events, the sIPSCs were acquired continuously in 2-minute sweeps during the entire recording time. The sIPSCs were recorded for 10 min in aCSF, followed by 12 min bath application of OT and followed by the drug washout (at least 10 more min after the OT application was completed). As sIPSCs are a combination of spike-independent IPSCs and spike-driven IPSCs, in some experiments the action potential generation was blocked with Tetrodotoxin (TTX, 1 μM, #1078; Tocris Bioscience, UK), leaving only the spike-independent IPSCs (miniature IPSCs, mIPSCs). GABA_A_ receptor antagonist, picrotoxin (25 μM), was used at the end of the recordings to verify the GABA-ergic origin of the sIPSCs. Finally, to study the effect of OT on the evoked inhibitory synaptic transmission in Type I-III neurons, we recorded the eIPSCs by electrical stimulation of the Stria Terminalis.

#### 2.3.5. CeA-projecting BNST_DL_ neurons

To determine an electrophysiological phenotype of BNST→CeA neurons, rats were first bilaterally injected into the CeA with a retrograde tracer, green microbeads (Lumafluor Inc., 100 nl; from Bregma: AP: -2.8 mm, ML: +4.0 mm, DV: -7.9 mm), and were allowed to recover for 4-7 days before electrophysiological recordings. Stereotaxic surgeries were performed as before (Martinon et al., 2019; Moaddab and Dabrowska, 2017). The slices were illuminated with CoolLED pE-300lite (Scientifica Ltd., Uckfield, UK) and ET - EGFP (FITC/Cy2) filter (Chroma Technology Corp, Bellows Falls, VT, US) to enable visually guided patching of fluorescent neurons. In a subset of these experiments we recorded sIPSCs from the fluorescent BNST→CeA neurons after application of OT as above. Slices were stored in 10% stabilized formalin post-recordings, mounted with Krystalon Mounting medium (Millipore Sigma, #64969-71), and analyzed with Nikon Eclipse E600 microscope (Melville, NY, US) and Olympus Fv10i confocal microscope for high magnification imaging (Center Valley, PA, US). Brain sections containing the CeA were also analyzed for the microbeads injection site. CeA was visualized with immunofluorescence using primary rabbit anti-enkephalin antibody (RA14124, Neuromics, MS, US) and secondary anti-rabbit Alexa Fluor 568 antibody (Life technologies Corporation, NY, US) according to the protocol as before (Dabrowska et al., 2013b).

### 2.4. Statistical analysis

Data are presented as MEAN ± standard error of mean (SEM). Data in line graphs are shown as the individual values from all recorded neurons (gray lines) as well as MEAN values (thick black line). Distribution of data sets was first evaluated with Kolmogorov-Smirnov test. Based on the normal distribution, the effects of OT on the sIPSCs, excitability, and firing frequency were analyzed by a repeated measures (RM) one-way analysis of variance (ANOVA) or mixed effects model, or paired *t*-tests, when applicable. Where the F-ratio was significant, all-pairwise post hoc comparisons were made using Tukey’s or Sidak’s tests. The statistical analyses were completed using GraphPad Prism version 9.0.0 (GraphPad Software Inc., San Diego, CA). *P* < 0.05 was considered significant.

## 3. Results

### 3.1. The effects of OT on intrinsic membrane properties and spontaneous firing in Type I-III BNST_DL_ neurons

In Type I neurons, OT induced a significant depolarization of RMP from -62.39 ± 0.84 mV to -57.94 ± 1.39 mV (*P* = 0.0002, F (1.593, 19.12) = 15.76, n = 14, mixed-effect analysis) in comparison to baseline (pre, *P* = 0.0013) and washout (post, *P* = 0.0304, **Fig. 1A-B**). This was associated with a significant increase of Rin from 170.98 ± 8.6 MΩ to 233.64 ± 18.6 MΩ (*P* = 0.0085, n = 9, paired *t*-test, **Fig. 1C**). As a consequence, OT induced a significant reduction of Rh from 21.96 ± 2.29 pA to 17.32 ± 2.26 pA (*P* = 0.0457, F (1.341, 16.09) = 4.226, n = 14, mixed-effect analysis) in comparison to baseline (pre, *P* = 0.0023, **Fig. 1D**). The Th of the first spike was also significantly reduced from -38.38 ± 0.68 mV to -39.75 ± 0.88 mV after OT (*P* = 0.0164, F (1.472, 20.61) = 5.749), in comparison to baseline (pre, *P* = 0.0346, **Fig. 1E**). When the OTR antagonist (OTA) was applied for 10 min before and during OT application, OT did not significantly affect RMP (*P* = 0.0710), Rh (*P* = 0.1231), Th (*P* = 0.3532) or Lat (*P* = 0.6252, **not shown**). In contrast to Type I neurons, OT did not affect the intrinsic membrane properties of Type II and III neurons (**Table 1**).

**Table 1.** Intrinsic membrane properties and firing activity of Type I-III BNST_DL_ neurons after OT (0.2 µM) application respective to baseline (pre-OT). \*\*\*\**P* < 0.0001, \*\**P* < 0.01, \**P* < 0.05. In Type II neurons OT did not affect RMP (*P* = 0.0985), Rin (*P* = 0.1377), Rh (*P* = 0.0964), Th (*P* = 0.5027), mAHP (*P* = 0.9147) or lAHP (*P* = 0.6340). Similarly, in Type III neurons OT did not significantly change RMP (*P* = 0.1066), Rin (*P* = 0.0788), Rh (*P* = 0.1664), Th (*P* = 0.5150), mAHP (*P* = 0.0775) or lAHP (*P* = 0.6472).

| **Type I neurons** | **Pre** | **OT** |
| --- | --- | --- |
| RMP (mV) | -62.39 ± 0.84 | **57.94 ± 1.39 \*\*** |
| Rin (MOhms) | 170.98 ± 8.6 | **233.64 ± 18.6 \*** |
| Rh (pA) | 21.96 ± 2.29 | **17.32 ± 2.26 \*\*** |
| Th (mV) | -38.38 ± 0.68 | **39.75 ± 0.88 \*** |
| Lat (ms) | 326.44 ± 28.63 | **224.58 ± 32.51 \*\*** |
| fAHP (mV) | 8.63 ± 0.89 | **7.47 ± 0.77 \*\*** |
| mAHP (mV) | 10.69 ± 0.61 | **8.45 ± 0.72 \*\*\*\*** |
| IAHP (mV) | 0.49 ± 0.12 | 0.41 ± 0.1 |
| **Type II neurons** |  |  |
| RMP (mV) | -57.04 ± 0.86 | -56.74 ± 0.78 |
| Rin (MOhms) | 221.8 ± 29.27 | 276.03 ± 46.66 |
| Rh (pA) | 16.92 ± 2.52 | 15 ± 2.66 |
| Th (mV) | -39.87 ± 0.95 | -40.32 ± 1.05 |
| mAHP (mV) | 10.96 ± 0.83 | 10.79 ± 0.86 |
| IAHP (mV) | 1.58 ± 0.33 | 1.41 ± 0.39 |
| **Type III neurons** |  |  |
| RMP (mV) | -65.73 ± 1.42 | -63.3 ± 1.83 |
| Rin (MOhms) | 120.66 ± 23.12 | 156.93 ± 29.7 |
| Rh (pA) | 56.46 ± 8.92 | 48.33 ± 7.36 |
| Th (mV) | -37.28 ± 0.85 | -37.08 ± 0.93 |
| mAHP (mV) | 8.68 ± 0.62 | 7.07 ± 0.93 |
| IAHP (mV) | 0.26 ± 0.12 | 0.22 ± 0.04 |

**Figure 1.**
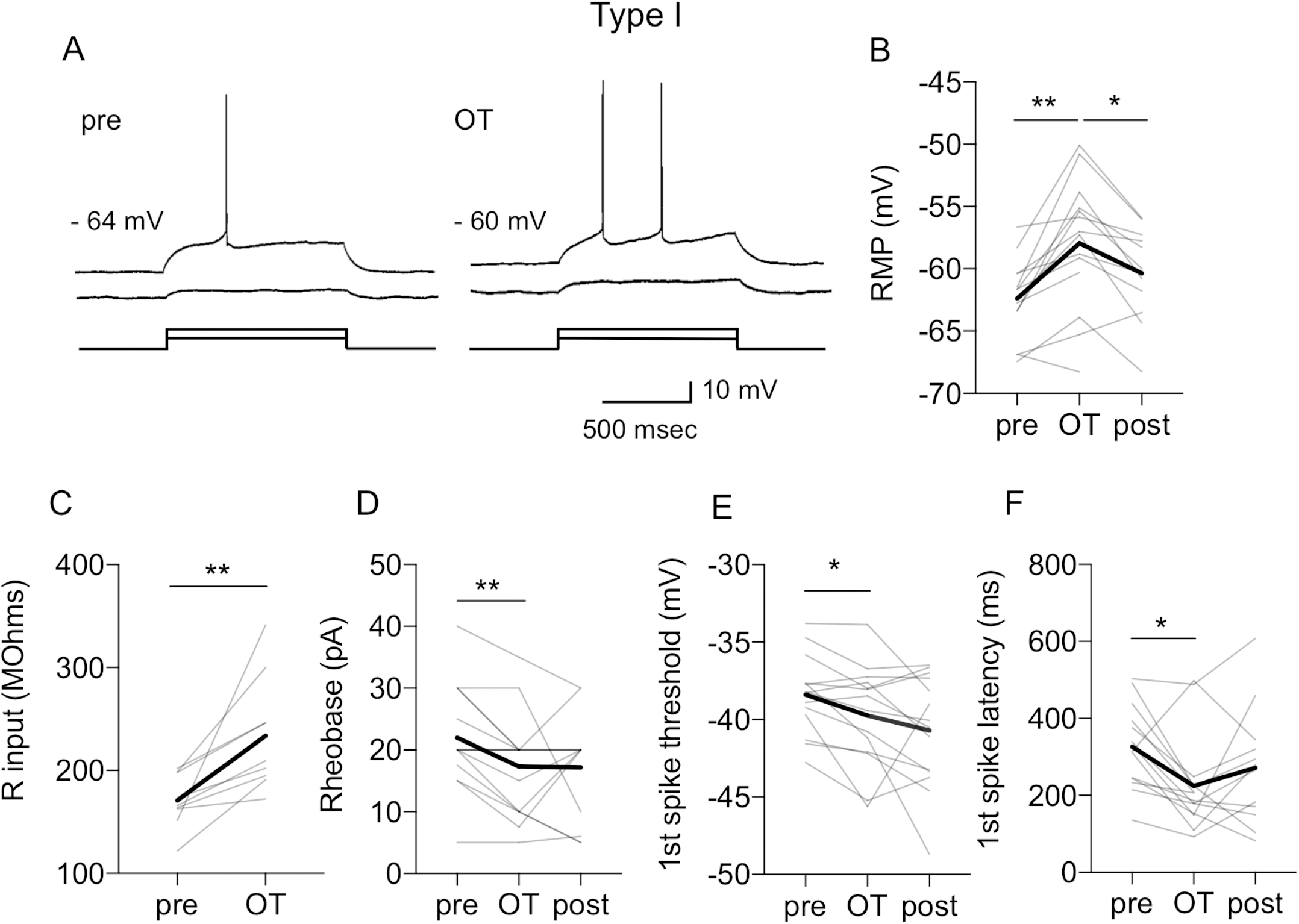
Oxytocin affects intrinsic membrane properties of Type I BNST_DL_ neurons. **A**: Representative traces showing that oxytocin (OT, 0.2 µM) shortens the latency to the 1^st^ spike at rheobase current (Rh) and depolarizes resting membrane potential (RMP, from -64 mV to -60 mV). This is associated with an increased input resistance (Rin), measured with a depolarizing current step below the Rh. OT also reduces Rh from 30 pA (pre) to 20 pA (OT) **B-F**: Line graphs with individual neurons responses (thin gray) and average responses for all neurons recorded (thick black) showing that OT significantly increases the RMP (**B**, *P* = 0.0002, F (1.593, 19.12) = 15.76, n = 14, mixed-effect analysis), increases Rin (**C**, *P* = 0.0085, n = 9, paired *t*-test), reduces the Rh (**D**, *P* = 0.0457, F (1.341, 16.09) = 4.226, n = 14, mixed-effect analysis), reduces the 1^st^ spike threshold (**E**, *P* = 0.0164, F (1.472, 20.61) = 5.749, and reduces the 1^st^ spike latency (**F**, *P* = 0.0185, F (1.971, 23.66) = 4.775, n = 14, mixed-effect analysis). Gray lines represent individual neurons’ responses to OT application. Thick black lines show average responses for all Type I neurons recorded, \*\**P* < 0.01, \**P* < 0.05 in comparison to pre or post-OT.

In addition, OT caused a leftward shift of input/output (I-O) relationship in Type I neurons, demonstrating a significant increment effect on SSF at 40 pA (*P* = 0.0024, F (1.667, 15.01) = 10.11, n = 12), at 50 pA (*P* = 0.0271, F (1.863, 16.76) = 4.628, n = 11), and at 60 pA (*P* = 0.0271, F (1.688, 11.82) = 5.267, mixed-effects analysis, n = 10). Post-hoc test showed a significant increase of the SSF after OT from 6.81 ± 0.68 Hz to 8.96 ± 0.63 Hz in comparison to baseline (pre, *P* = 0.0032) and washout (post, *P* = 0.0095) at 40 pA; from 8.73 ± 0.73 Hz to 11.20 ± 0.70 Hz (*P* = 0.0163, pre and *P* = 0.0226, post) at 50 pA; and from 9.22 ± 0.57 Hz to 11.89 ± 0.99 Hz (*P* = 0.0271, pre and *P* = 0.0054, post) at 60 pA. The OT effect reached a trend at 30 pA (from 5.64 ± 0.5 Hz to 7.31 ± 0.53 Hz, *P* = 0.0820) and at 70 pA (from 10.79 ± 0.6 Hz to 13.62 ± 0.99 Hz, *P* = 0.0794). The effect was not significant at 20 pA (*P* = 0.1268) and 80 pA (*P* = 0.1320, **Fig. 2A-B**). Importantly, in the presence of OTA, OT did not affect the SSF of Type I neurons at 80 pA (*P* = 0.7960), 70 pA (*P* = 0.4237), 60 pA (*P* = 0.6312), 50 pA (*P* = 0.4299), 40 pA (*P* = 0.3365), or 30 pA (*P* = 0.4817, **Fig. 2C**). OT did not affect SSF in Type II (F (1, 7) = 0.0020, *P* = 0.9652) or Type III neurons (F (1, 8) = 2.907, *P* = 0.1266, **Fig. 2D-E**).

**Figure 2.**
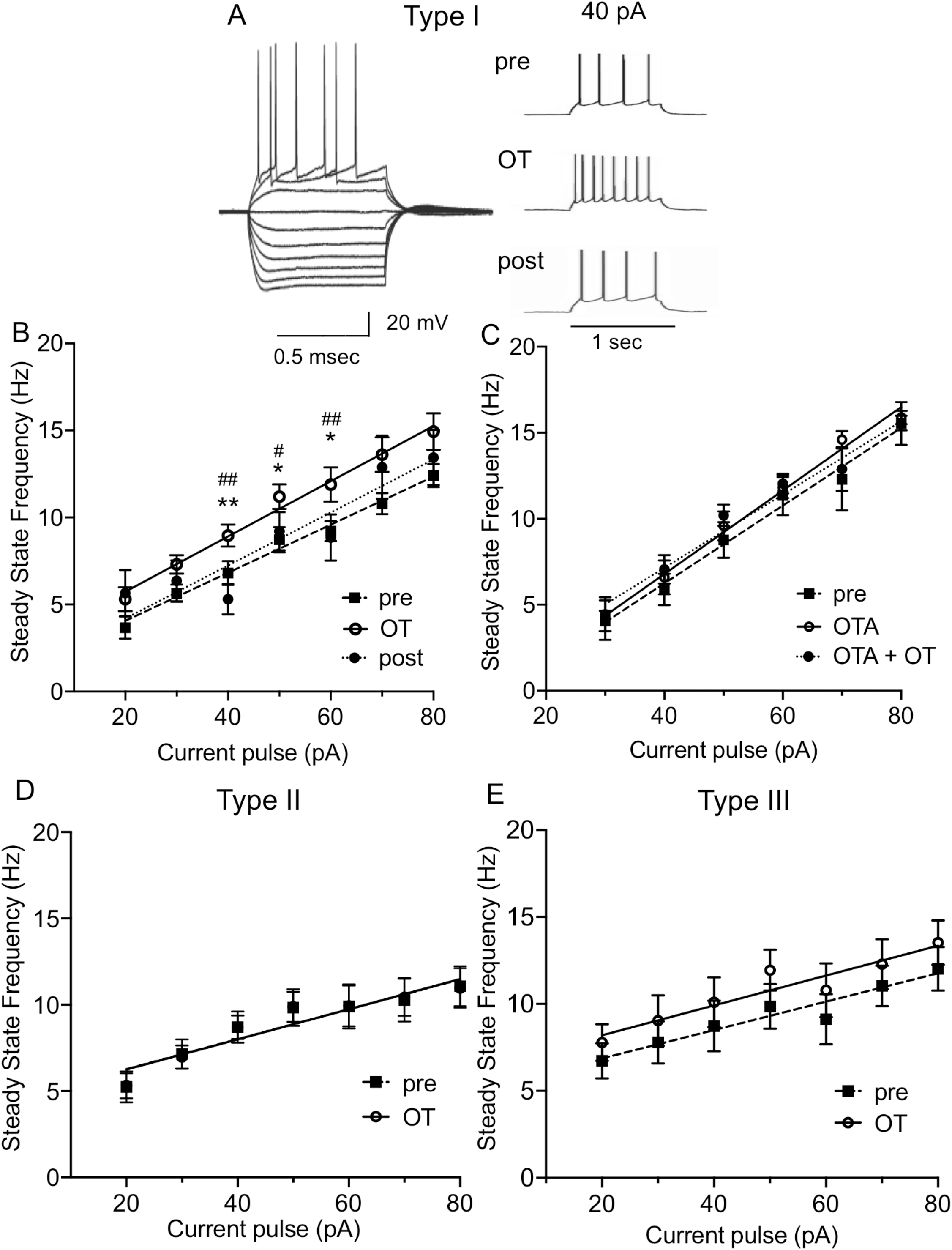
Oxytocin increases intrinsic excitability of Type I, but not Type II or Type III, BNST_DL_ neurons. **A**: Representative traces showing voltage deflections in response to hyperpolarizing and depolarizing current injections characteristic of Type I neuron (left) and the increase in firing frequency of Type I neuron in the presence of OT respective to baseline (pre) (right). **B**: OT causes a leftward shift of input/output (I-O) relationship in Type I neurons. The steady state frequency (SSF) was determined as an average of an inverse of the inter-spike intervals (ISI) from all action potentials starting from the second action potential. In Type I neurons, OT shows a significant increment effect on SSF at 40 pA (*P* = 0.0024, F (1.667, 15.01) = 10.11), at 50 pA (*P* = 0.0271, F (1.863, 16.76) = 4.628), and at 60 pA (*P* = 0.0271, F (1.688, 11.82) = 5.267, mixed-effects analysis). **C**: Application of OTR antagonist, OTA (0.4 µM), prevents the leftward shift of the I-O relationship induced by OT application (40 pA, *P* = 0.3365, 50 pA, *P* = 0.4299, 60 pA, *P* = 0.6312, n = 5, RM ANOVA). **D-E:** OT application does not modify the I-O relationship in Type II (**D**) or Type III BNST_DL_ neurons (**E**). Each data point represents the mean ± SEM, \*\**P* < 0.01, \**P* < 0.05 in comparison to pre-OT, ## *P* < 0.01, # *P* < 0.05 in comparison to post-OT.

This increase in the spiking ability of Type I neurons after OT was associated with a significant shortening of the first spike’s latency from 326.44 ± 28.63 msec to 224.58 ± 32.51 msec (*P* = 0.0185, F (1.971, 23.66) = 4.775, n = 14, mixed-effect analysis), in comparison to baseline (pre, *P* = 0.0083, **Fig. 1F**). We also investigated the effects of OT on the fAHP, the mAHP and the lAHP. In Type I neurons, OT significantly reduced the amplitude of fAHP (*P* = 0.0124, F (1.473, 17.68 = 6.493, mixed-effects analysis, n = 14), from 8.63 ± 0.89 mV to 7.47 ± 0.77 mV, *P* = 0.0062, in comparison to pre, **Fig. 3A-B**) and mAHP (*P* <0.0001, F (1.844, 22.13) = 23.25, mixed effect analysis, n = 14), from 10.69 ± 0.61 mV to 8.45 ± 0.72 mV (*P* = 0.0001 vs. pre, and *P* = 0.0003 vs. post), **Fig. 3A, D**). Notably, OT did not affect the amplitudes of fAHP (*P* = 0.2220, n = 7, **Fig. 3C**) and mAHP (*P* = 0.3968, n = 7, RM one way-ANOVA, **Fig. 3E**) in the presence of OTA. Lastly, OT did not significantly affect the amplitude of lAHP in Type I neurons (*P* = 0.2401, **Table 1**). In contrast to Type I neurons, OT did not affect the amplitude of mAHP and lAHP of Type II and III neurons (**Table 1**). We did not measure the effect of OT on fAHP in these neurons, because the fAHP is not present in Type II and Type III neurons.

**Figure 3.**
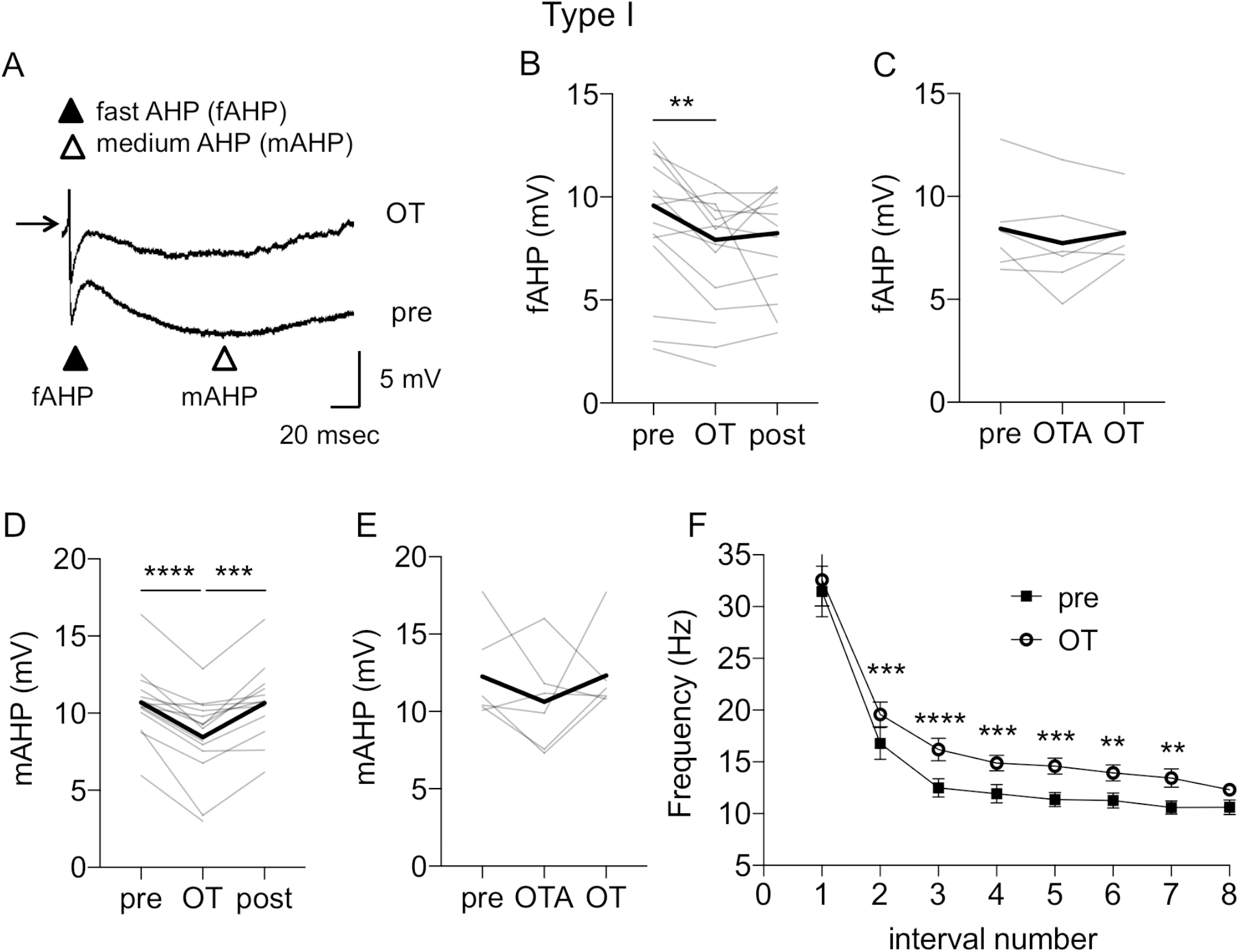
Oxytocin reduces fast and medium afterhyperpolarizations and modifies early and late spike frequency adaptation in Type I BNST_DL_ neurons. **A**: Representative traces from Type I neuron showing a reduction of fast AHP (fAHP) and medium AHP (mAHP) after OT application respective to baseline (pre). The amplitudes of the fAHP and mAHP following the first action potential evoked at Rh were calculated as the difference from the action potential threshold to the membrane voltage measured after 2-5 msec and 50-100 msec, respectively. The lAHP was measured at 2000 msec after the end of the depolarizing pulse, following generation of five action potentials. **B**: Line graphs showing individual neurons’ responses (gray) and reduction of the fAHP during OT application (*P* = 0.0124, F (1.473, 17.68 = 6.493, mixed-effects analysis, n = 14). **C**: OTR antagonist, OTA (0.4 µM), prevents the OT-induced reduction of fAHP (*P* = 0.2220, n = 7, RM RM ANOVA). **D**: Line graph showing reduction of mAHP after OT application (*P* <0.0001, F (1.844, 22.13) = 23.25, mixed effect analysis, n = 14). **E**: OT-induced reduction of mAHP is blocked by OTR antagonist, OTA (*P* = 0.3968, n = 7, RM ANOVA). Thick black lines show average responses for all Type I neurons recorded. **F**: OT modifies early and late spike frequency adaptation. The early and late spike frequency adaptation was investigated using a depolarizing pulse of 1 sec and measuring the instantaneous frequency for every action potential interval. The ISI number is the inter-stimulus interval between two consecutive action potentials. OT significantly increased the instantaneous frequency of ISI (*P* < 0.0001, F (1, 13) = 82.69, mixed effects model), **B-E**: Gray lines represent individual neurons’ responses to OT application. Thick black lines show average responses for all Type I neurons recorded. **F**: Each data point represents the MEAN ± SEM, \*\*\*\**P* < 0.0001, \*\*\**P* < 0.001, \*\**P* < 0.01.

We also investigated if these changes in AHP amplitude could modify the spike frequency adaptation. We found that OT significantly increased the instantaneous frequency of ISIs (*P* < 0.0001, F (1, 13) = 82.69, n = 13, mixed effects model) and post-hoc test showed a significant OT effect on frequency (in Hz) of ISI number 2 (ISI-2) from 16.77 ± 1.55 to 19.57 ± 1.2 (*P* = 0.0006), ISI-3 from 12.48 ± 0.88 to 16.19 ± 1.09 (*P* < 0.0001), ISI-4 from 11.92 ± 0.88 to 14.87 ± 0.76 (*P* = 0.0006), ISI-5 from 11.36 ± 0.68 to 14.58 ± 0.77 (*P* = 0.0001), ISI-6 from 11.26 ± 0.72 to 13.93 ± 0.76 (*P* = 0.0079), and ISI-7 from 10.58 ± 0.64 to 13.44 ± 0.88 (*P* = 0.0051, **Fig. 3F**).

During cell-attached recordings, the spontaneous firing was present in 7 out of 9 Type I neurons and OT significantly increased the firing rate in the spontaneously firing neurons from 0.88 ± 0.66 Hz to 2.19 ± 0.93 Hz (*P* = 0.0242, paired *t-*test, **Fig. 4A-B**). In addition, although only 7 out of 19 recorded Type II neurons showed spontaneous firing in a cell-attached mode, in all spontaneously firing Type II neurons, OT significantly reduced the firing rate from 1.96 ± 0.63 Hz to 1.29 ± 0.60 Hz, (*P* = 0.0304, paired *t*-test, n = 7, **Fig. 4C-D**). In addition, in Type II neurons recorded with high seal (>2GΩ), which did not fire spontaneously, we also estimated the effect of OT on the RMP by measuring the voltage in cell-attached current clamp mode as described before (Perkins, 2006) and we show that OT application does not change the RMP (pre 58.6 ± 2.52 mV to 58.5 ± 2.42 mV, n = 6).

**Figure 4.**
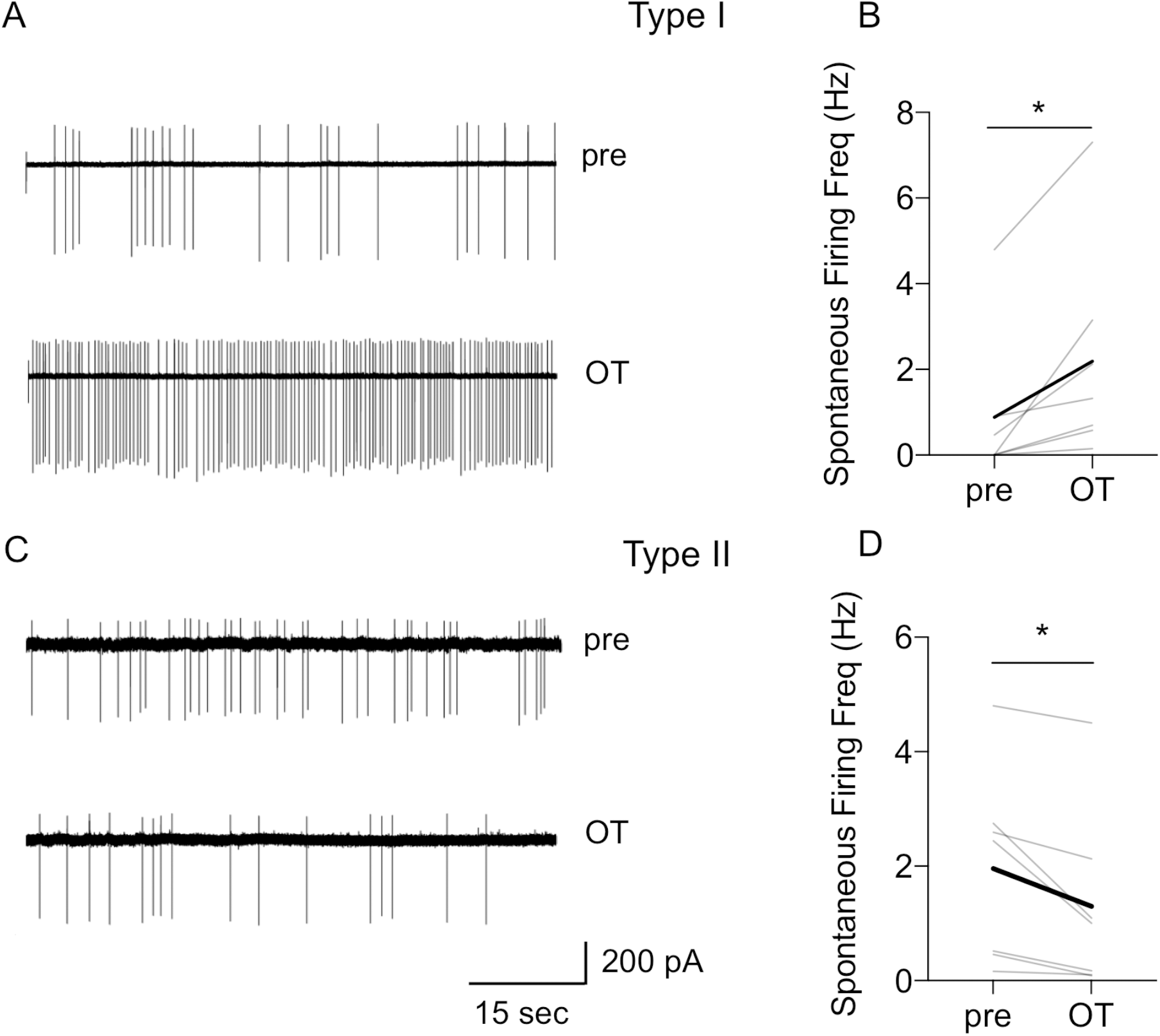
Oxytocin increases spontaneous firing rate of Type I neurons and reduces spontaneous firing rate of Type II BNST_DL_ neurons in cell-attached recordings. **A**: Representative traces of Type I neuron showing an increase in spontaneous firing rate after OT application. **B**: Line graph showing that OT increased spontaneous firing rate in all spontaneously firing Type I neurons (*P* = 0.0242, paired *t-*test, n = 7). **C**: Representative traces of Type II neuron showing a reduction in spontaneous firing rate after OT application. **D**: Line graph showing that OT significantly reduced spontaneous firing rate in spontaneously firing Type II neurons in cell-attached mode (*P* = 0.0304, paired *t*-test, n = 7), gray lines represent individual neurons’ responses, thick black line represents average response, \**P* < 0.05.

Overall, these results suggest that OT increases the intrinsic excitability and facilitates the spontaneous firing selectively in Type I BNST_DL_ neurons, whereas it reduces spontaneous firing rate in Type II BNST_DL_ neurons.

### 3.2. The effects of OT on inhibitory synaptic transmission in Type I-III BNST_DL_ neurons

In Type I neurons, OT did not affect the sIPSCs frequency (*P* = 0.3018), amplitude (*P* = 0.3347), or total charge (*P* = 0.6087, n = 7, paired *t-*test, **not shown**). Moreover, OT did not change the amplitude of the eIPSC in Type I neurons (*P* = 0.2449, paired *t*-test, n = 5, **not shown**).

In Type II BNST_DL_ neurons, application of OT caused a significant increase in the frequency of sIPSCs in 12 out of 14 recorded neurons (*P* = 0.0022, F (1.058, 12.17) = 14.41, n = 14, mixed-effect analysis) from 1.03 ± 0.20 Hz to 2.64 ± 0.47 Hz vs. baseline (pre, *P* = 0.0033), and vs. washout (post, *P* = 0.0018, **Fig. 5A-B**). Notably, the effect of OT on the sIPSCs frequency was not associated with a significant change in the amplitude (*P* = 0.5855, F (1.563, 17.19) = 0.4718, n = 14, mixed-effect analysis, **Fig. 5C**) or in the total charge (*P* = 0.5744, **not shown**). OT did not increase the sIPSCs frequency in the presence of OTA (*P* = 0.1788, F (1.469, 8.814) = 2.135, n = 6, RM ANOVA, **Fig. 5D**). In addition, when the spike-driven sIPSCs were blocked with TTX (*P* = 0.0151, F (1.030, 6.179) = 10.98, n = 6, RM ANOVA), OT no longer affected the frequency of miniature IPSCs (mIPSCs) of Type II neurons (*P* = 0.9975, **Fig. 5E**). Finally, OT did not affect the eIPSCs amplitude (at the Vh of +10 mV) in Type II neurons, from 266.76 ± 55.43 pA (pre) to 316.17 ± 82.49 pA (post, *P* = 0.2459, paired *t*-test) (**Fig. 6A** and **6C**). These results demonstrate that OT indirectly increases the frequency of spike-driven sIPSCs in Type II neurons.

**Figure 5.**
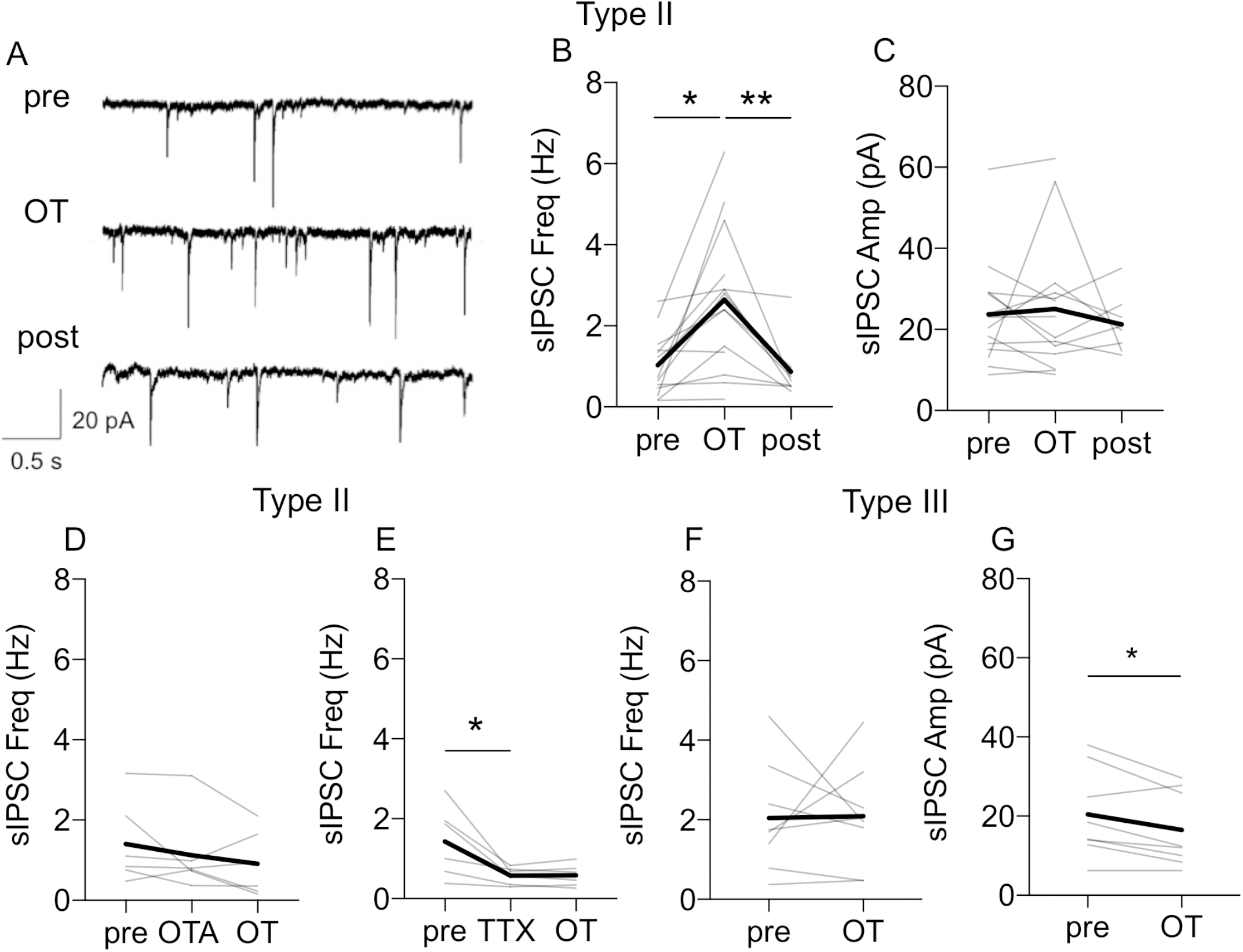
Oxytocin modulates inhibitory synaptic transmission in BNST_DL_ neurons. **A**: Representative recordings from Type II neuron showing an increase in the frequency of spontaneous inhibitory postsynaptic currents (sIPSCs) after OT application respective to baseline (pre). **B-C**: Line graphs (gray) showing that 12 out of 14 recorded Type II neurons showed an increase in sIPSCs frequency (**B**, *P* = 0.0022, F (1.058, 12.17) = 14.41, n = 14, mixed-effect analysis) but not amplitude (**C**, *P* = 0.5855, F (1.563, 17.19) = 0.4718, n = 14, mixed-effect analysis) after OT application. **D**: OT does not affect sIPSCs frequency in a presence of OTR antagonist, OTA (0.4 µM) (*P* = 0.1788, F (1.469, 8.814) = 2.135, n = 6, RM ANOVA). **E**: When spike-driven IPSCs were blocked with TTX (*P* = 0.0151, F (1.030, 6.179) = 10.98, n = 6, RM ANOVA), OT does not affect the sIPSCs frequency. **F-G**: OT does not affect frequency (**F**, *P* = 0.9362) but reduces amplitude of sIPSCs in Type III neurons (**G**, *P* = 0.0160, paired *t*-test. Gray lines represent individual neurons’ responses to OT application. Thick black lines show average responses for all Type I neurons recorded, \*\**P* < 0.01, \**P* < 0.05 in comparison to pre- or post-OT.

**Figure 6.**
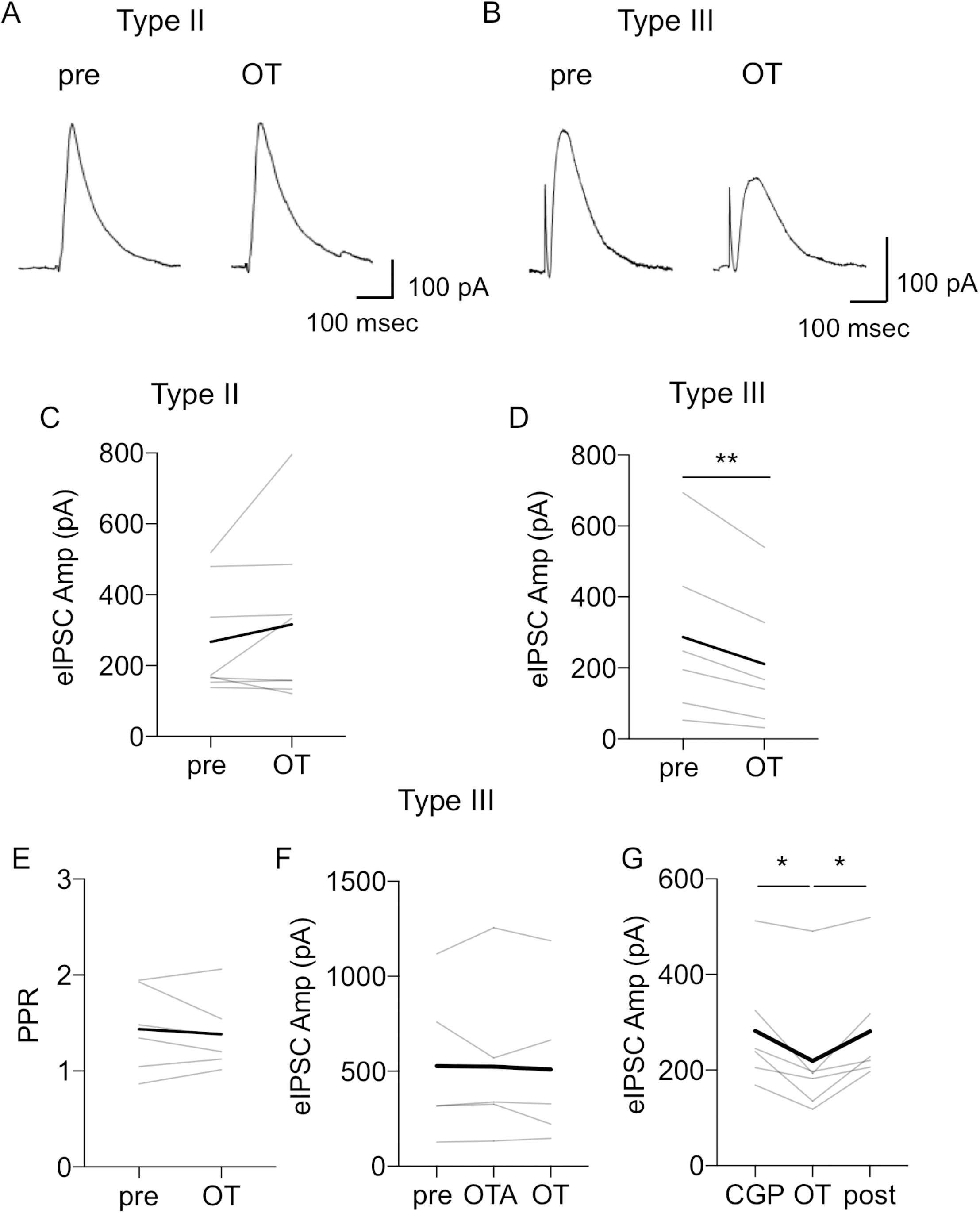
Oxytocin reduces the amplitude of evoked IPSCs in Type III, but not Type II neurons. **A-B**: Representative traces of eIPSCs in response to electrical stimulation of the stria terminalis in Type II (**A**) and Type III (**B**) neurons. **C**: Line graph showing that OT does not change the mean amplitude of eIPSC in Type II neurons (*P* = 0.2459, paired *t*-test, n = 5). Thick black line shows averaged responses for all Type II neurons recorded. **D-G**: Line graphs showing responses of individual Type III neurons to OT application and the reduction of eIPSC amplitude (**D**, *P* = 0.0105, paired *t*-test, n = 6), without any significant change in the PPR (**E**, *P* = 0.5672). This effect of OT on eIPSCs amplitude was blocked by OTR antagonist, OTA (**F**, F (1.421, 7.105) = 0.1722, *P* = 0.7739, RM ANOVA, n = 6), but not by GABA-B receptor antagonist, CGP 55845 (**G**, F (1.258, 6.291) = 11.16, *P* = 0.0122, RM ANOVA, n = 6). Line graphs (gray lines) represent individual neurons’ responses to treatment. Thick black lines show average responses for all neurons recorded.

In Type III neurons, OT did not affect the frequency (*P* = 0.9440, **Fig. 5F**), but it reduced amplitude (*P* = 0.0314, **Fig. 5G**) and total charge of sIPSCs (*P* = 0.0372, paired *t*-test, n = 6, **not shown**). In addition, in Type III neurons, OT significantly reduced the amplitude of the eIPSCs from 286.76 ± 97.41 pA (pre) to 210.69 ± 78.57 pA (*P* = 0.0105, paired *t*-test, n = 6, **Fig. 6B** and **6D**). This reduction of the eIPSC amplitude was not associated with a significant change in the PPR (*P* = 0.5672, paired *t*-test, **Fig. 6E**). Notably, in a presence of OTR antagonist, OT did not reduce the amplitude of eIPSCs in Type III neurons (F (1.421, 7.105) = 0.1722, *P* = 0.7739, RM ANOVA, n = 6), **Fig. 6F**). However, OT significantly reduced the amplitude of the eIPSCs in the presence of the GABA-B receptor antagonist CGP 55845 from 282.45 ± 50.58 pA to 219.38 ± 55.83 pA (F (1.258, 6.291) = 11.16, *P* = 0.0122, RM ANOVA, n = 6, **Fig. 6G**), showing a significant reduction in comparison to pre *(P* = 0.0413), and post (*P* = 0.0374). Again, this reduction of the eIPSC amplitude after OT in the presence of CGP 55845 was not associated with a significant change in the PPR (F (1.438, 7.191) = 2.560, *P* = 0.1499, RM ANOVA, n = 6, **not shown**). Overall, these findings suggest that in Type III neurons OT affects the synaptic response through a postsynaptic mechanism.

### 3.3. CeA-projecting BNST_DL_ neurons

In rats injected with microbeads into the CeA (**Fig. 7A**), retro-labeled fluorescent neurons were found in subdivisions of the BNST_DL_, primarily in the oval nucleus of the BNST_DL_ (BNST_OV_) and to a lesser extent in anteromedial BNST (BNST_AM_, **Fig. 7B-C**). We recorded 13 fluorescent neurons from 11 rats injected with microbeads to the CeA (**Fig. 7D**), and all 13-recorded neurons were classified as Type II (**Fig. 7E**). To determine OT effects in these Type II BNST→CeA neurons, we performed whole cell patch-clamp recordings in 5 of these Type II fluorescent neurons and we measured frequency and amplitude of sIPSCs before and after OT application. We demonstrate that OT increased the sIPSCs frequency (F (1.004, 4.016) = 10.99, *P* = 0.0293, RM ANOVA, n = 5) from 1.64 ± 0.41 Hz to 3.95 ± 0.52 Hz (vs. pre, *P* = 0.0416 (**Fig. 7F**), but not amplitude (F (1.007, 4.026) = 3.099, *P* = 0.1527, **not shown**) in the fluorescent Type II BNST→CeA neurons, showing that OT inhibits the Type II BNST→CeA output neurons (**Fig. 7G**).

**Figure 7.**
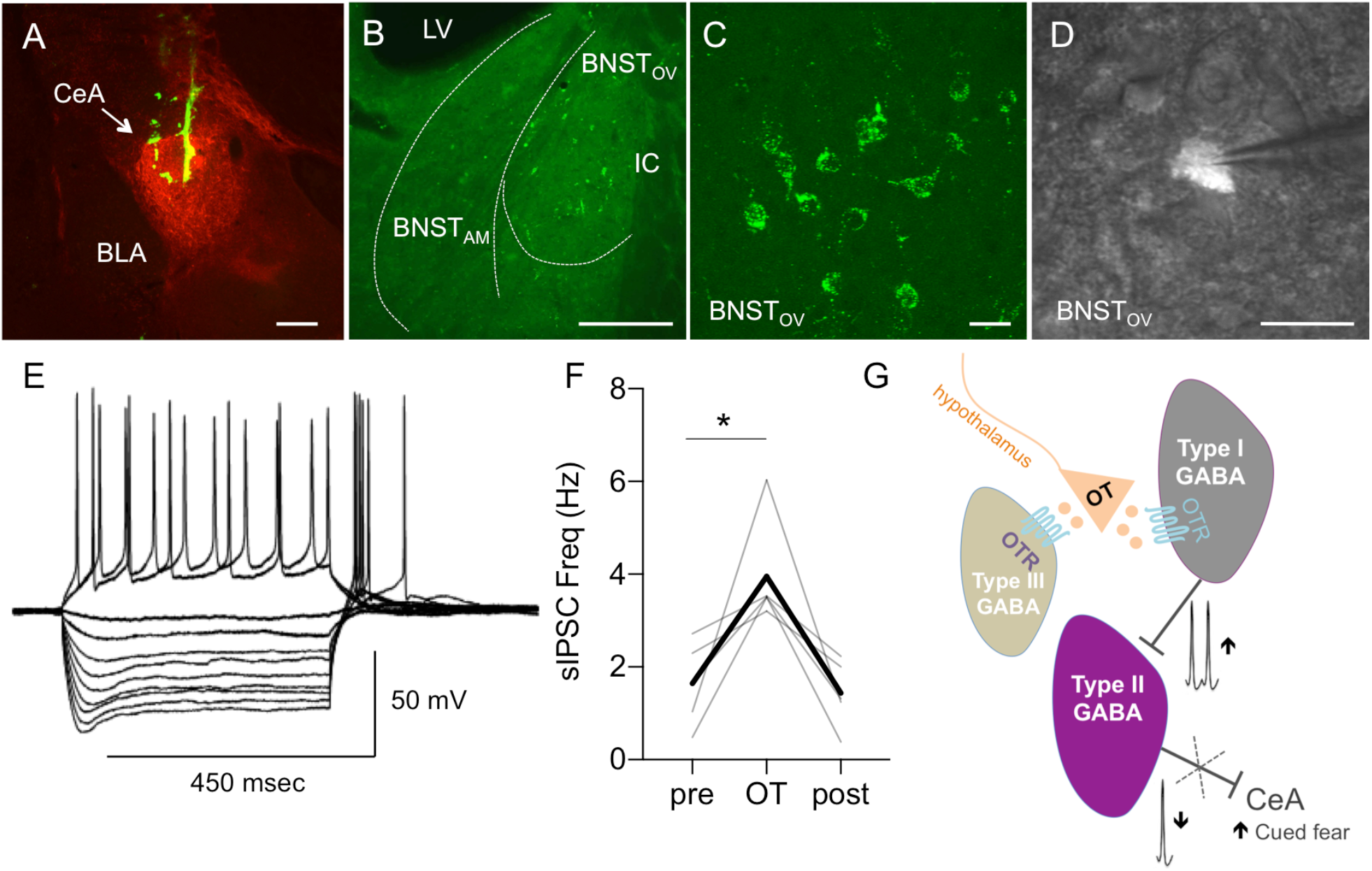
Oxytocin inhibits Type II BNST→ CeA output neurons. **A-B**: Microphotographs showing a representative injection site of green retrobeads in the central amygdala (CeA, here visualized with enkephalin immunofluorescence in red, **A**, scale bar 100 µm) and neurons expressing green fluorescent microbeads in the BNST_DL_ (**B**, bar 100 µm). BNST neurons projecting to the CeA (BNST→CeA neurons) are found in the dorsolateral BNST, primarily the oval nucleus (BNST_OV_), and to a lesser extent in the anteromedial BNST (BNST_AM_) (BLA – basolateral nucleus of the amygdala, IC – Internal capsule, LV – Lateral ventricle). **C**: High magnification (60x) microphotographs obtained from a confocal microscope showing green fluorescent beads present in membranes and processes of the BNST→CeA neurons (scale bar 10 µm). **D-E**: Electrophysiological characterization of the BNST→CeA neurons using visually guided patching (**D**, 40x, scale bar 10 µm). Representative recording of the BNST→CeA fluorescent neuron showing the electrophysiological properties of a Type II neuron such as the presence of a sag during the hyperpolarized current pulses and the spike rebound at the end of the 450 msec current pulse injection (**E**). **F**: OT increases the frequency of sIPSCs (F (1.004, 4.016) = 10.99, *P* = 0.0293, RM ANOVA, n = 5) in the fluorescent Type II BNST→CeA neurons. Gray lines represent individual neurons’ responses to treatment. Thick black lines show average responses for all Type II neurons recorded. **G**: Diagram showing that OT regulates the intrinsic inhibitory network of the BNST_DL_. By directly increasing spontaneous firing of Type I regular spiking interneurons, OT reduces firing of Type II BNST_DL_ neurons, which send projections to the central amygdala (BNST→CeA neurons). OT also reduces the evoked GABA-ergic inhibition in Type III BNST_DL_ neurons, via a direct postsynaptic mechanism.

## 4. Discussion

Here we demonstrate that OT regulates the intrinsic inhibitory network of the BNST_DL_, where it increases intrinsic excitability and spontaneous firing of Type I, regular spiking interneurons, which leads to a potentiation of GABA-ergic inhibition and a reduction of spontaneous firing of Type II, burst firing neurons. Using visually guided patching of retro-labeled BNST_DL_ neurons, we confirm that Type II are output neurons of the BNST_DL_, which project to the CeA (BNST→CeA) and we show that OT inhibits these Type II BNST→CeA neurons. Furthermore, we show that OT reduces the spontaneous and evoked GABA-ergic inhibition in Type III BNST_DL_ neurons, via a postsynaptic mechanism. Therefore, we demonstrate that there are three groups of OT-responsive neurons in the BNST_DL_; a population of interneurons (Type I) that is directly excited by OTR activation, a group of BNST→CeA projection neurons (Type II), which are indirectly inhibited by OT, and a group of Type III neurons, which demonstrate reduced GABA-ergic inhibition after OT application (see **graphical abstract and Fig. 7G**). The OT-mediated inhibition of the BNST→CeA output neurons has powerful implications for the modulation of fear and anxiety-like behaviors.

Neurons of the BNST_DL_ are GABA-ergic (Cullinan et al., 1993; Dabrowska et al., 2013a; Sun and Cassell, 1993), and can be categorized into three major types (I-III) (Daniel et al., 2017; Hammack et al., 2007). Type I neurons, characterized by regular firing and lack of burst-firing activity (Hammack et al., 2007; Hazra et al., 2011), were classified as interneurons by Paré and colleagues (Rodríguez-Sierra et al., 2013). This has been recently supported by retrograde labeling showing a very small fraction of Type I neurons (in comparison to Type II and Type III) projecting outside of the BNST_DL_ (Yamauchi et al., 2018). We show that Type I neurons are the primary OT-responsive neurons of the BNST_DL_ as demonstrated by their increased excitability and spontaneous firing rate after OT application. Previous studies using single unit recordings showed that OT increased the spontaneous firing rate of 50% of BNST neurons in lactating female rats (Ingram et al., 1990), and the magnitude of the response was significantly potentiated during lactation (Ingram and Moos, 1992). Notably, in our study, the excitatory effects of OT in Type I neurons are paralleled by increased inhibitory transmission observed in Type II BNST_DL_ neurons. The specific effect of OT on the sIPSCs frequency, but not amplitude, suggests a presynaptic mechanism in Type II neurons. In addition, since the effect was no longer observed when the spike-driven IPSCs were suppressed with TTX, this indicates an indirect influence driven by an increased spontaneous firing of BNST_DL_ interneurons. This has been further supported by the lack of the OT effect on eIPSCs in Type II neurons, suggesting that the inhibitory effect of OT in Type II neurons is not a result of the stimulation of the extrinsic inhibitory projections via stria terminalis.

In contrast, in Type III neurons, OT reduces the amplitude and the total charge, but not the frequency of sIPSCs and eIPSCs, without changing the PPR. The effect of OT on eIPSCs was also observed in the presence of an antagonist for presynaptic GABA-B receptors, further excluding presynaptic effect. Overall, these results indicate a postsynaptic mechanism of OT in Type III neurons. OT was shown previously to reduce evoked GABA-ergic transmission in the CeA (Tunstall et al., 2019) and to depress GABA-ergic synapses in the supraoptic nucleus of the hypothalamus via postsynaptic increase of Ca^2+^ due to the activation of G-protein coupled receptors (Alger and Nicoll, 1980; Brussaard et al., 1996). However, since in our hands the effect on the amplitude was accompanied by a reduction in the total charge, this could potentially indicate shortening of GABA channel opening (Martina et al., 1994) via OTR-mediated increases in intracellular Ca^2+^, as demonstrated after OTR activation before (Brussaard et al., 1996). Therefore, in Type III neurons OT might reduce the amplitude of sIPSCs and eIPSC through a postsynaptic mechanism involving signal transduction-mediated regulation of intracellular Ca^2+^ levels.

To further investigate the OT effects on intrinsic excitability and firing, we demonstrated that OT reduced fAHP and mAHP selectively in Type I neurons. AHPs generated after a single action potential or a train of action potentials limit the frequency of neuronal firing (Alger and Nicoll, 1980), as demonstrated by a significant increase in action potential firing after pharmacological inhibition of AHPs (Pedarzani and Storm, 1993). The fAHP, mediated by BK channels, is responsible for early spike frequency adaptation in many neuronal types (Gu et al., 2007). The mAHP is mediated by the non-inactivating M-current, which blockade with selective antagonist reduces late spike adaptation (Gu et al., 2008, 2005b). An OTR-mediated increase of neuronal excitability was recently shown in the hippocampus, where a selective OTR agonist induced membrane depolarization and increased membrane resistance in CA2 pyramidal cells leading to a leftward shift in their firing frequency-current relationship (Tirko et al., 2018). Here we show that OT-mediated excitation of Type I neurons in the BNST_DL_ was associated with a significant increase in instantaneous frequency of action potentials due to the reduction in the early and late adaptation. Therefore, by acting on potassium conductances OT might increase the intrinsic excitability and the spontaneous firing of Type I neurons.

According to our knowledge, this is the first study utilizing cell-attached mode in a cell-type specific manner to show that in the BNST_DL_, the majority of Type I neurons fire spontaneously, and this effect is further potentiated by OT. Notably, although we showed that the majority of Type II neurons remain silent even during cell-attached configuration, in all spontaneously firing Type II neurons, OT application reduced the rate of firing. Notably, the indirect inhibitory effect on spontaneous firing of Type II neurons was further confirmed by the lack of the OT effect on current-evoked firing in these neurons, which was investigated in a presence of synaptic transmission blockers. The latter further suggests that OT does not alter intrinsic properties of Type II neurons and instead, reduces the spontaneous firing rate and so their inhibitory synaptic transmission via the direct excitatory effect on Type I neurons.

The BNST was not entirely isolated during recordings, such as the slice also potentially contained GABA-ergic neurons in the medial preoptic area (Okamura et al., 1990), which express OTR (Sharma et al., 2019), and project to the BNST (Simerly and Swanson, 1988). However, this projection primarily targets the principal encapsulated nucleus in the posterior BNST (Maejima et al., 2015), where OTR are involved in reproductive behavior (Kremarik et al., 1991). Notably, the posterior BNST was not a part of our slice preparation. In addition, although we cannot entirely rule out a possibility that some inhibitory inputs to the BNST_DL_ might be originating from anteromedial (BNST_AM_) or ventral BNST (BNST_VEN_), which were included in our slice, the intrinsic inhibitory projections in the BNST have been primarily shown to originate in the BNST_DL_, mainly the BNST_OV_, and target BNST_AM_ and BNST_VEN_, whereas opposite projections are rare (Gungor and Paré, 2016). Therefore, our results strongly suggest that Type I neurons are BNST_DL_ interneurons providing inhibitory synaptic input to Type II neurons, an effect potentiated by OT.

It is important to note that the BNST also contains neurons producing arginine-vasopressin (AVP) (al-Shamma and De Vries, 1996) and has a high expression of AVP receptors, V1AR (Grundwald et al., 2016). As OT shows a relatively high affinity toward V1AR, this raises a possibility of non-specific effects via V1AR. However, all the significant OT effects in our study were blocked by OTA, which is highly selective toward OTR with a negligible affinity toward V1AR, see (Manning et al., 2012).

Similar modulation of the intrinsic inhibitory network by OT was shown in a variety of limbic brain regions, including the CeA. The CeA and the BNST share many similarities, including predominantly GABA-ergic nature, analogous peptidergic cell types, as well as similar intrinsic and extrinsic connectivity, and together they form an extended amygdala complex (Alheid, 2003; Sun and Cassell, 1993). Interestingly, in the CeA, there are two groups of OT-responsive neurons. A group of interneurons located in the lateral nucleus of the CeA (CeL) (Gozzi et al., 2010) is excited by OTR activation (Huber et al., 2005; Terburg et al., 2018; Viviani et al., 2011), which strengthens inhibitory input to medial nucleus of the CeA (CeM) output neurons, which were shown to projecting to lateral periaqueductal gray and reduce contextual fear (Viviani et al., 2011), for review see (Beyeler and Dabrowska, 2020). Similarly to the effects we observed in the BNST_DL_, OTR-mediated indirect inhibition of the CeM output neurons was demonstrated as increased frequency of sIPSC during whole-cell recordings and reduced spontaneous firing rate in a cell-attached configuration. Similarly, in the hippocampus (Harden and Frazier, 2016; Maniezzi et al., 2019; Owen et al., 2013; Tirko et al., 2018; Zaninetti and Raggenbass, 2000), lateral amygdala (Crane et al., 2020), and the prefrontal cortex (Marlin et al., 2015; Nakajima et al., 2014), OT was shown to excite a population of interneurons, including fast-spiking parvalbumin interneurons, which were shown to inhibit glutamatergic projection neurons.

We show that in the BNST_DL_ OT inhibits Type II neurons, whereas it reduces the evoked inhibition on Type III neurons. Recently, both Type II and Type III were demonstrated output neurons of the BNST_DL_, with major projections to the CeA (62% of Type II and 33% of Type III), LH (83% of Type III), and the VTA (17% of Type II, 79% of Type III neurons). In our experiments, all the retrogradely labeled BNST→CeA neurons we recorded from the BNST_OV_ were identified as Type II neurons, and OT increased the frequency of sIPSCs in these Type II BNST→CeA neurons. Since all the BNST→CeA neurons we recorded from were identified as Type II, it is possible that Type II vs. Type III neurons target different subdivisions of the CeA as the BNST→CeA neurons were shown projecting to the CeL and the CeM (Gungor et al., 2015. It would still need to be determined which Type III projection neurons are affected by OT. Although neuropeptidergic phenotypes corresponding to Type I-III neurons remain elusive, Type III are putative corticotropin-releasing factor (CRF) neurons of the BNST_OV_ (Dabrowska et al., 2013b), which were shown projecting to the LH, VTA, and the CeA, among many other brain regions (Dabrowska et al., 2016). Therefore, by reducing GABA-ergic transmission in Type III neurons, OT might increase probability of firing of these Type III/CRF projection neurons. However, as some CRF neurons in the BNST_DL_ were also shown to serve as local interneurons (Partridge et al., 2016), OT might have an excitatory influence on not just one (Type I) but possibly on two different types of BNST_DL_ interneurons (Type I and Type III/CRF neurons). As a part of the future studies we will determine whether OT reduces GABA-ergic transmission in Type III interneurons vs. Type III projection neurons.

Notably, Type II neurons are the most abundant and heterogeneous neuronal population in rat BNST_DL_ (Hazra et al., 2011; Yamauchi et al., 2018). The BNST is known to mediate stress-induced anxiety-like responses (Dabrowska et al., 2013b; Daniel and Rainnie, 2016; Davis et al., 2010; Sparta et al., 2013), contextual fear (Davis et al., 2010; Sullivan et al., 2004; Waddell et al., 2006), as well as fear responses to un-signaled, diffuse, or unpredictable threats (Duvarci et al., 2009; Goode et al., 2020, 2019), but might also inhibit cued fear responses to predictable threats (Meloni et al., 2006). Although the projection from the BNST_DL_ to the CeA has been shown before (McDonald et al., 1999), a functional inhibitory projection was recently demonstrated (Gungor et al., 2015; Yamauchi et al., 2018), and was shown to facilitate anxiety-like behavior (Yamauchi et al., 2018). In apparent contrast to the BNST-mediated anxiety, our recent behavioral findings demonstrate that OTR transmission in the BNST_DL_ reduces fear responses to un-signaled/diffuse threats and facilitates acquisition of cued fear (Janeček and Dabrowska, 2019; Martinon et al., 2019; Moaddab and Dabrowska, 2017). Based on the well-established role of the CeA in the acquisition of cued fear (Wilensky et al., 2006), our current electrophysiology results suggest that OTR activation in the BNST_DL_ facilitates cued fear and reduces anxiety by inhibiting the Type II BNST→CeA output neurons. Therefore, our findings are of great significance for understanding the role of the BNST and its interactions with the CeA in the neurobiology of fear and anxiety and further highlight the unique potential of targeting oxytocin receptors for the treatment of anxiety disorders (Marvar et al., 2021).

## 5. Acknowledgements

We would like to thank Dr. Attilio Caravelli for his help with Matlab. We also thank Rachel Chudoba for proofreading the manuscript. This work was supported by grant from the National Institute of Mental Health R01MH113007 and R01MH113007-04S1 to JD and start-up funds from the Chicago Medical School, Rosalind Franklin University of Medicine and Science to JD.

## 6. Contributions

JD initiated the study. JD and WF designed the study. WF, FB, and VOP acquired data. WF, FB, VOP, and JD analyzed data. WF, FB, VOP and JD interpreted data. JD and WF wrote the manuscript. JD obtained funding.

